# Whole genome analysis of ExPEC ST73 from a single hospital over a 2-year period identified different circulating clonal groups

**DOI:** 10.1101/428599

**Authors:** DR Bogema, J McKinnon, M Liu, N Hitchick, N Miller, C Venturini, J Iredell, AE Darling, P Roy Chowdury, SP Djordjevic

**Affiliations:** Elizabeth Macarthur Agricultural Institute, NSW Department of Primary Industries, Menangle, NSW, 2568 Australia; The ithree Institute, University of Technology Sydney, NSW, 2007 Australia; San Pathology, Sydney Adventist Hospital, Wahroonga, NSW, 2076 Australia; Centre for Infectious Diseases and Microbiology, The Westmead Institute for Medical Research, The University of Sydney, Westmead, NSW, 2145 Australia; Westmead Hospital, Westmead, NSW, 2145 Australia

**Keywords:** Uropathogenic Escherichia coli, class 1 integron

## Abstract

ST73 has emerged as one of the most frequently isolated extraintestinal pathogenic *E. coli* (ExPEC). To examine the localised diversity of ST73 clonal groups including their mobile genetic elements profile, we sequenced the genomes of 16 multiple drug-resistant ST73 isolates from patients with urinary tract infection from a single hospital in Sydney, Australia between 2009 and 2011. Genome sequences were used to generate a SNP-based phylogenetic tree to determine the relationship of these isolates in a global context with ST73 sequences (n=210) from public databases. There was no evidence of a dominant outbreak strain of ST73 in patients from this hospital, rather we identified at least eight separate groups, several of which reoccur, over a two-year period. The inferred phylogeny of all ST73 strains (n=226) including the ST73 Clone D i2 reference genome shows high bootstrap support and clusters into four major groups which correlate with serotype. The Sydney ST73 strains carry a wide variety of virulence-associated genes but the presence of *iss*, *pic* and several iron acquisition operons was notable.

**Impact:** ST73 is a major clonal lineage of ExPEC that causes urinary tract infections often with uroseptic sequelae but has not garnered substantial scientific interest as the globally disseminated ST131. Isolation of multiple antimicrobial resistant variants of ExPEC ST73 have increased in frequency, but little is known about the carriage of class 1 integrons in this sequence type and the plasmids that are likely to mobilise them. This pilot study examines the ST73 isolates within a single hospital in Sydney Australia and provides the first large-scale core-genome phylogenetic analysis of ST73 utilizing public sequence read datasets. We used this analysis to identify at least 8 sub-groups of ST73 within this single hospital. Mobile genetic elements associated with antibiotic resistance were less diverse and only three class 1 integron structures were identified, all sharing the same basic structure suggesting that the acquisition of drug resistance is a recent event. Genomic epidemiological studies are needed to further characterise established and emerging clonal populations of multiple drug resistant ExPEC to identify sources and aid outbreak investigations.

## Introduction

Extraintestinal pathogenic *Escherichia coli* (ExPEC) are phylogenetically diverse and comprise uropathogenic *E. coli* (UPEC), neonatal meningitis-causing *E. coli* (NMEC) and avian pathogenic *E. coli* (APEC). ExPEC account for ~75–95% of urinary tract infections (UTI). A proportion of these infections can spread from the urinary tract with invasion of epithelial cells in the bladder (cystitis) and kidney cells (pyelonephritis), and transmission to systemic circulation (blood sepsis), posing a serious threat to human health. ExPEC are enteric bacteria but their capacity to capture a wide array of virulence-associated genes (VAGs) by lateral gene transfer has expanded the repertoire of niches they colonise. ExPEC may carry diverse and often redundant combinations of VAGs whose impact on human health remains ill-defined. Epidemiological studies indicate that a subset of pathogenic *E. coli* lineages including ST73, ST131, ST405, ST393, ST69, ST95, ST10, ST38, and ST127 [1–5] are responsible for most ExPEC infections [3, 6–8]. Carriage of combinations of virulence genes enhance virulence [9], however, carriage of antimicrobial resistance genes, particularly those encoding ESBLs and fluoroquinolones, as well as an ability to cause opportunistic infections in vulnerable (elderly) hosts, may also contribute to virulence. It is notable that none of these hypotheses have been experimentally validated [3, 6].

ExPEC have become the leading cause of blood sepsis in Europe [10]. Notable in this regard is the alarming rise in the incidence of ST73, now one of the most frequently isolated UPEC globally and the leading cause of bacteraemia in the East Midlands region of the UK [1, 11–14]. ST73 belongs to Clermont phylogroup B2 and is known to display different serogroups with serotype O6 predominating (ST73-O6-B2) [15, 16]. It has recently been suggested that the rise in the incidence of multiple drug resistant (MDR) ST73 in the UK is not due to the emergence of a dominant clone because they are genetically diverse and carry a different array of plasmids encoding resistance to multiple antimicrobials. Many recently described isolates of ST73 carry genes that encode extended spectrum β-lactamases and resistance to antimicrobials used in veterinary medicine [14]. This seems to be a recent adaptation in this ST as previously characterized ST73 isolates from cases of uncomplicated UTI sourced from Greece, Portugal, Sweden and the UK, resulted susceptible to most clinically relevant antimicrobials [17] and most (75%) did not carry plasmids, classic vehicles of MDR [17]. These data combined with the most recent findings seem to suggest that the rise in the carriage of MDR plasmids in ST73 may be a recent concerning event [14, 18].

Here we have characterised whole genome sequences of 15 class 1 integrase (*intI1*)-containing ST73 strains from a hospital in Sydney. To determine if these highly localised strains were from a limited number of clonal lineages, phylogenetic inferences were made by comparing SNP differences in core genomes shared by ST73 strains from our Sydney collection with those from six high-quality reference genome sequences and ST73 strains (n=204) from seven countries sourced from global sequence read archives. We also examined mobile and chromosomal genetic content within this localised isolate cohort to further examine their diversity. We compiled the repertoire of antimicrobial genes and virulence genes and mapped the class 1 integron structures carried by these isolates. As carriage of the class 1 integrase is considered a reliable proxy for multiple drug resistance [19], we used S1-PFGE followed by Southern hybridization with an *intI1* probe to examine plasmid content and carriage of the class 1 integrase on plasmids.

## Methods

### Isolate source and culture conditions

Clinical samples in this project were from a larger collection obtained from Sydney Adventist Hospital from 2009–2011. Bacterial species were identified by the VITEK^®^ 2 (bioMérieux) system at Sydney Adventist Hospital. For DNA extraction, strains were first grown on a Lysogeny Broth (LB) agar plate to isolate single colonies of which one was used to inoculate 2 mL of LB broth followed by shaking for 16 hours at 37°C. Antibiotic susceptibility testing for ampicillin, cefotaxime, chloramphenicol, streptomycin and sulfafurazole was performed via the CDS method [20]. These antibiotics were selected based on antibiotic resistance gene content inferred from genome sequencing.

### Nucleic acid purification and whole genome sequencing

*E. coli* DNA was extracted using the Isolate II Genomic DNA Extraction kit (Bioline) according to the manufacturer’s instructions. For each sample, tagmentation of genomic DNA, and PCR amplification of tagged DNA were performed in triplicate using the Nextera system (Illumina). Sequencing libraries were pooled, then cleaned and size selected using SPRI beads (Beckman Coulter). Normalization was guided by read counts obtained from a Nano flowcell run on a MiSeq instrument. Agilent 2100 Bioanalyzer, with High Sensitivity DNA kit was used to quantitate the pooled library before loading onto an Illumina HiSeq. Paired-end 150 bp reads were generated using the HiSeq 2500 v4 system.

### Genome assembly and gene presence

Genome assembly was achieved with raw reads using the A5-miseq pipeline [21] and checked for consistency by additional assembly with SPAdes 3.9.0 [22]. Antibiotic resistance and virulence-associated genes were identified from assembled genomes using BLASTn and SRST2 [23]. Searches were performed against antibiotic resistance genes sourced from the ARG-ANNOT V3 database and a panel of virulence-associated genes identified from the Virulence Factors[24] Database (VFDB) and literature searches [25]. Serotyping was performed *in silico* with SRST2 using EcOH sequences supplied with this package. Draft genome reads obtained from SRA were searched for a minimal set of marker genes derived from integron structures characterised in the Sydney strains. Low-quality alignments based on SRST2 output were discounted (n=3).

### Archived sequence read selection

All additional ST73 sequences not generated by this study were obtained from complete whole genome assemblies (n = 6) [26–30] and public sequence read archives (NCBI-EMBL). Raw Illumina reads sourced from ST73 isolates (n = 284) were considered for SNP-based phylogenetic analysis, including strains sequenced in this study from the Sydney Adventist Hospital (n = 16), isolates with host, source and isolation location meta-data identified from the EnteroBase database (n = 246; http://enterobase.warwick.ac.uk/; accessed 5/12/2016) and a previous ST73-focused study from the United Kingdom (n = 22) [11]. Samples were excluded if sequence type could not be confirmed as ST73 using SRST2 (n = 4). Further samples were excluded (n = 30) if isolate status could not be confirmed by BioProject meta-data or where the description of methods could not be identified by an associated publication [31–38]. Samples were additionally excluded (n = 30) if they produced low reference genome coverage (>90%) in whole-genome alignments. Additional sample filtering of ST73 reads is described in Table S2.

### S1-PFGE analysis

The complement of large (>20 kb) plasmids in each bacterial isolate was determined by S1 nuclease (Promega, Madison, WI, USA) digestion and pulsed-field gel electrophoresis (PFGE) as described previously [39, 40]. Southern blot hybridization was used to determine the genomic location of the *intI1* gene. PCR amplicons for *intI1* were obtained using published primers (int1F and int1R; [41]) and labelled using the PCR DIG Probe Synthesis Kit (Roche, Mannheim, Germany). DNA was transferred from the S1-PFGE gel to a nylon membrane (GE Health, Little Chalfont, UK) using a VacuAid vacuum transfer apparatus (HybAid, Teddington, UK) and hybridisation was performed using the Roche DIG Filter Hybridization system (Roche, Mannheim, Germany) following manufacturer’s instructions. Images were acquired on a ChemiDoc™ MP System (Bio-Rad Laboratories, Richmond, CA, USA).

### SNP based phylogenetic analyses

Our initial attempts to examine ST73 phylogeny with our 16 genome sequences and 6 complete whole genome sequences (CFT073, ABU83972, ATCC25922, Nissle 1917, clone D i2 and clone D i14) using marker gene approaches [42] provided limited resolving power (Fig. S1). Consequently, to more appropriately examine these ST73 strains, we employed SNP-based phylogenetic methods.

For SNP-based phylogenetic trees, core genome alignments were generated with Snippy v3.1.0 (https://github.com/tseemann/snippy) using default options. Briefly, reads were mapped using bwa mem v0.7 to an ST73 reference genome. Raw alignments were processed with samtools v1.3.1 and variants called using freebayes v1.1.0. SNP derived genomes were reconstructed using vcftools v0.1.14, with low-coverage (< 10 X) and degenerate reference positions filtered. Recombinant regions were removed using Gubbins v2.20 (options -i 10) [43] yielding aligned, SNP-derived, recombination-filtered core genomes. From this alignment, core genome phylogenetic trees were inferred by maximum-likelihood using RAxML v8.2.9 [44]. Branch support was estimated by bootstrap analysis employing 100 replicate trees. Trees were rooted using the ultrametric tree method included with RAxML. Scripts used for the analysis of SNP phylogeny can be found online at https://github.com/bogemad/snp_phylogeny.

Initially we performed this analysis using only isolates sequenced in this study and a high-quality published ST73 reference (Fig. 1). To identify the most suitable reference genome for this purpose, we aligned reads individually to the six complete reference genomes (above) and examined reference sequence quality, core alignment lengths and final tree support values. Using this methodology, the most suitable reference was identified as clone D i2 which with archived public sequence reads generated a core genome of 3,818,344 bp, representing 75.8% of the ST73 clone D i2 sequence.

**Figure 1.**
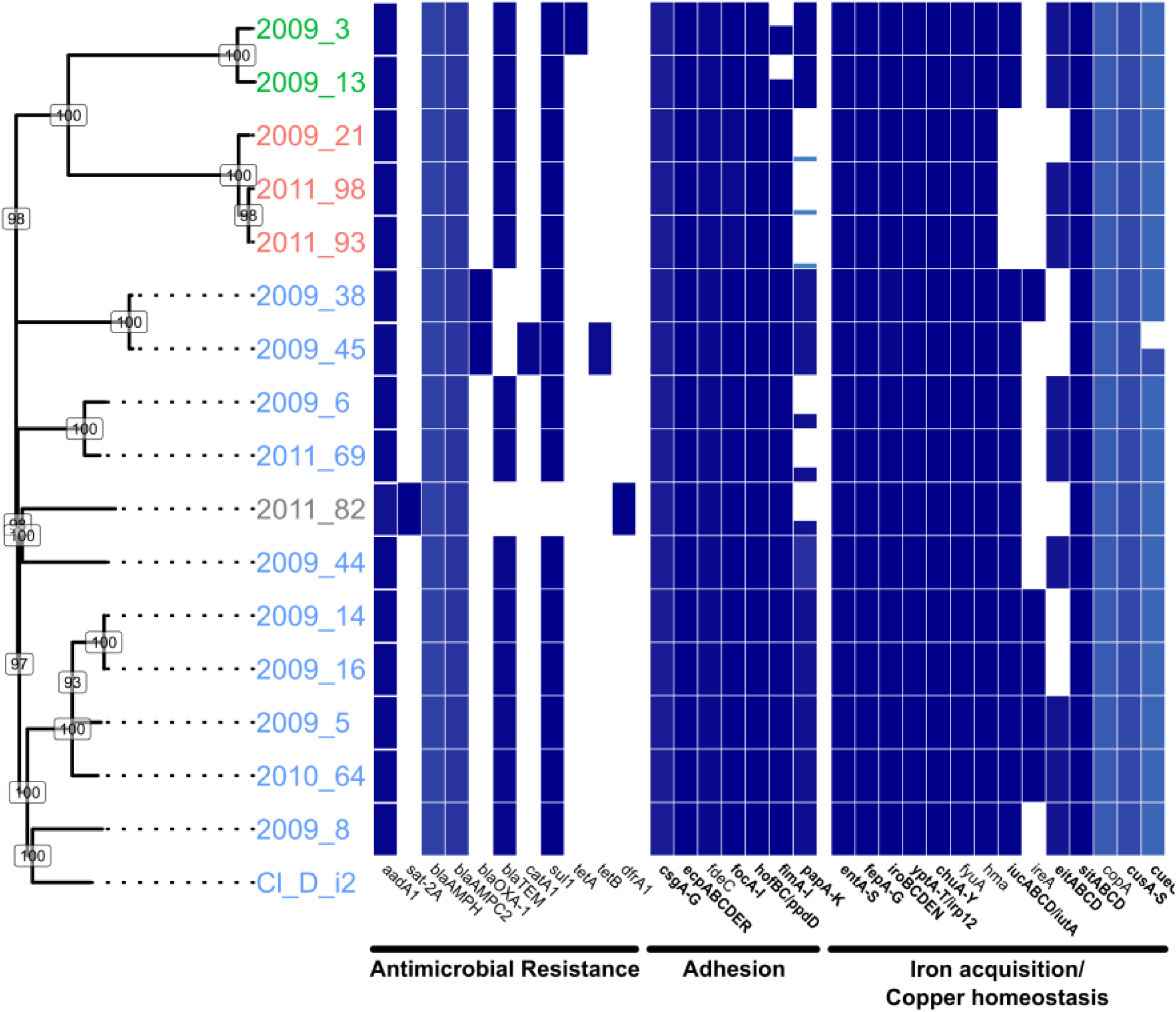
A SNP-dervied phylogenetic tree of the Sydney ST73 strains sequenced in this study compared with antimicrobial resistance and virulence profiles. Bootstrap values based on 100 replicate trees are shown at labeled nodes. Isolate serotypes as determined by *in silico* serotyping are shown by coloured tip labels. Antibiotic resistance and virulence gene/gene-family presence (blue) or absence (white) is shown by the linked bar-graph/heatmap. Percentage identity of BLAST matches is indicated by heatmap shade with darker shades representing higher identity. BLAST match coverage is represented by tile height with solid tiles representing 100% coverage. For gene families (x-axis; bold) tile height represents total BLAST match coverage of all gene family members and shows completeness of the gene family. Where single genes are indicated (x-axis; plain text), bar height represents BLAST match coverage of the gene.

## Results

### Assembly information and statistics

The genome sequences of 16 ST73 strains from the Sydney Adventist Hospital were determined here. The Whole Genome Shotgun project has been deposited at DDBJ/ENA/GenBank and the sequence read archive. Assembly statistics as well as accession numbers, number of sequencing reads, and amount of sequencing data used to generate assemblies can be found in Table S1.

### Public read high-throughput sequencing analysis

The Sydney isolates separated in several groups that closely aligned with *E. coli* serotype including O22:H1, O25:H1 and O6:H1. However, most of the Sydney isolates clustered within a larger O6:H1 group. To further interrogate observed diversity within the O6:H1 group of isolates and to place Sydney strains within a broader global context, we expanded the SNP-based phylogenetic analysis to include additional ST73 sequence reads obtained from public sequence read archives (NCBI-EMBL-DDBJ) and the aforementioned six complete ExPEC genomes with the 16 Sydney genomes. The inferred phylogeny of the 226 strains (Fig. 2; see Fig. S2 for all strain labels and branch support values) shows strong major branch support. Analysis of the SNP-derived phylogenetic tree shows correlation with observed serotypes O6:H1, O25:H1 and O22:H1, with most strains observed within the O6:H1 cluster. Strain 2011_82 could not be assigned an O-type from *in* silico serotyping but was identified as H1 and clustered most closely with O6:H1 isolates (Fig. 2 – Serotype).

**Figure 2.**
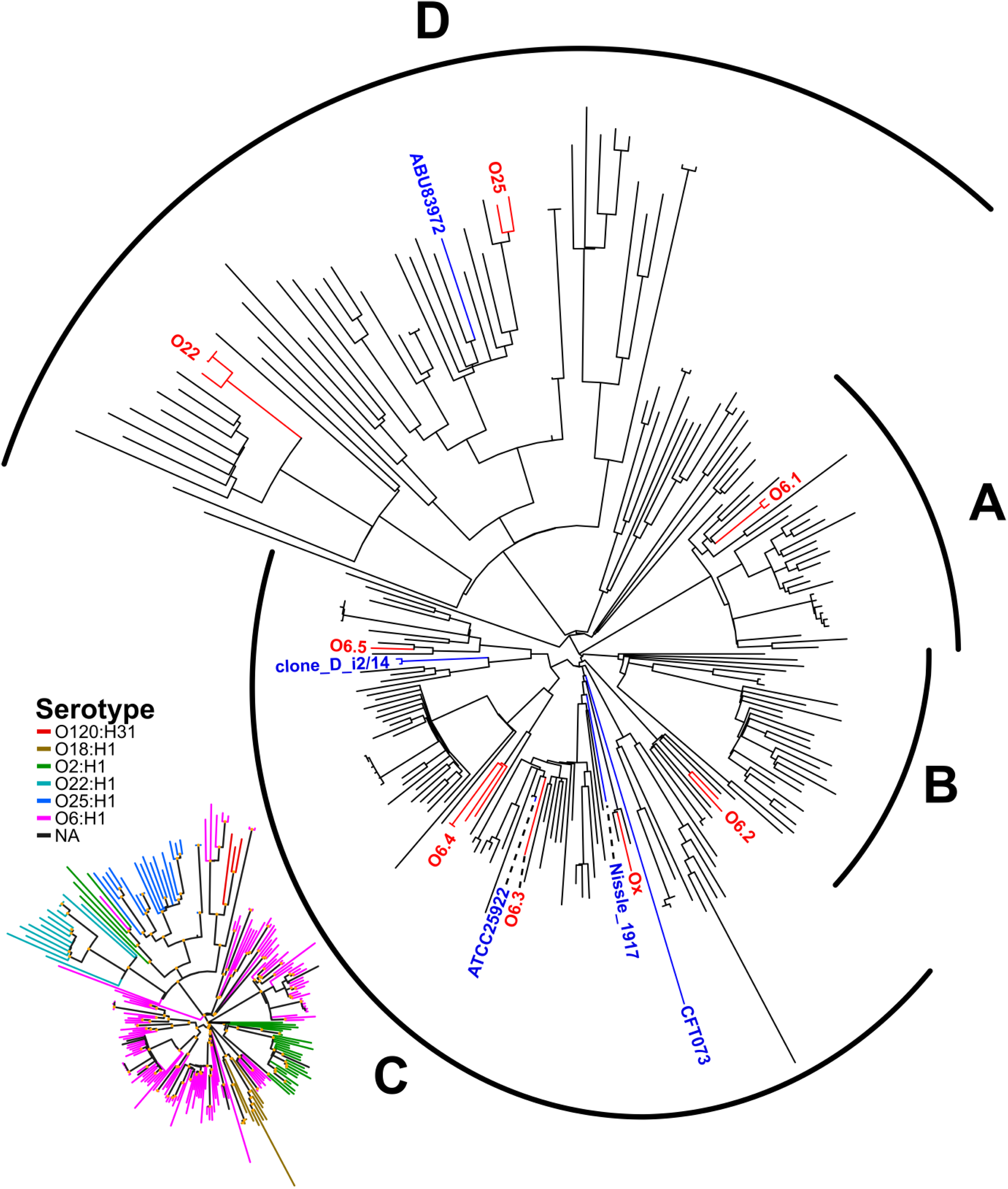
SNP-based maximum-likelihood phylogram of 226 ST73 strains. A more detailed tree with branch support values and tip labels can be found in Figure S2. ST73 isolates separate into 4 distinct groups, labelled A-D, which correlate well with *in silico* serotyping (insert). ST73 isolates sequenced in this study cluster into 8 distinct groups, shown in red, high-quality complete ST73 genomes are shown in blue. Trees were constructed using 18,426 SNPs identified by read mapping to the clone D i2 reference sequence, reduced from 27,568 SNPs by filtering of recombination regions.

Examination of ST73 phylogenetic structure reveals four significant clades (Fig. 2 and S2): one exclusively associated with serotype O6:H1 (Group A), one exclusively associated with serotype O2:H1 (Group B), one primarily serotype O6:H1 with a subclade of O18:H1 (Group C), and finally a polyphyletic group consisting of O2:H1, O6:H1, O22:H1, O25:H1 and O120:H31 serotypes (Group D). Strains sequenced from Sydney separated into eight distinct groups correlated with serotype (Fig. 2 – red labels). Isolates with O22:H1 and O25:H1 serotypes formed their own clusters, while serotype O6:H1 separated into 5 distinct clusters (O6-1: 2009_38/45; O6-2: 2009_6/2011_69; O6-3: 2009_44; O6-4: 2009_5/14/16/2010_64; O6-5:2009_8). Strain 2011_82 clustered with other O6:H1 strains and formed the final group (Ox).

### Virulence profiles of Sydney strains

We identified differences in virulence gene profiles of *E. coli* strains examined in this study (Fig. 1). Significantly, we found that virulence gene profiles were largely consistent in strains from the same phylogenetic groups O25, O22, O6-1 – O6-5, Ox (Fig. 1, see Table S4 and S5 for greater detail). For adhesion-related genes, critical components of P fimbriae (*papACDEFGHJK*) were absent in six strains (Fig. 1). Additionally, in strains of group O25, genes encoding the Type I fimbriae major subunit (*fimA*), periplasmic chaperone (*fimC*), regulatory subunit (*fimE*) and the fimbriae-associated *fimI* were missing in blastn searches. Other genes encoding F1C fimbriae, Curli fibres, Type IV pili, *E. coli* common pili and the *fdeC* adhesion genes were present in all strains. The importance of FdeC as a putative virulence factor is underpinned by the observation that it is i) a broadly conserved, *E. coli* adhesin whose expression is upregulated on the surface of UPEC when it contacts host cells; and ii) a major target during humoral immune responses that significantly reduced kidney colonization in mice challenged transurethrally with UPEC strain 536 [45].

Iron acquisition is critical for the growth of ExPEC in low iron environments *in vivo* and it is not uncommon to identify genes linked to siderophore production and processing in UPEC. Complete Enterobactin, Salmochelin, Yersiniabactin gene clusters were identified in all ST73 strains while Aerobactin genes were identified in all strains except those belonging to group O22. Genes for heme uptake including the *chu* operon and *hma* gene were present in all strains, as were those related to iron uptake such as the *sit* ABC transporter operon and ferric Yersinia uptake (*fyuA*) gene. In contrast, the putative iron uptake gene cluster *eitABCD* and adhesion/iron-uptake gene *ireA* were only identified in a subset of strains. In addition to iron uptake, genes encoding copper resistance have also been linked to virulence [46] and antimicrobial resistance [47]. The *cus* system, encoding a four-component copper efflux pump, was present and complete in all strains. However, in strain 2009_45, *cueR*, an important regulator controlling copper detoxification and efflux *copA* and *cueO* genes, was not located in all searches.

Larger differences were observed in the presence of toxin genes. Strains from group O25 and strain 2009_8 contained the highest number of toxin genes including cytotoxic necrotizing factor 1 (*cnf1*), the hemolysin (hlyABCD) cluster, hemolysin E (*hlyE*) and secreted autotransporter toxin (*sat*). Genes that have been previously shown to promote propagation of *E. coli* in blood, such as proteases *pic* and *tsh* and the increased serum survival (*iss*) gene were present in all strains as well as cellular invasion promoting *ibe* gene cluster. Closely related *hek* and *tia* genes, associated with epithelial cell invasion in Neonatal Meningitis-causing and enterotoxigenic *E. coli* respectively, are both found in separate strains. Furthermore, *tcpC* associated with immune modulation via inhibition of Toll/interleukin-1 (IL-1) receptor signalling, was only found in groups O6-1–5 and Ox.

### Antibiotic resistance

All *intI1* positive isolates were tested for resistance to ampicillin, cefotaxime, chloramphenicol, streptomycin, sulfafurazole and trimethoprim using the CDS method (Table 1). Strain 2011_82 did not have a class 1 integrase gene and was not tested. All strains were resistant to ampicillin, streptomycin and sulfafurazole. Genes encoding resistance to these antibiotics were all accounted for in the genome sequence data by the class 1 integron-associated genes *aadA1* and *sul1* as well as one of three *bla* gene variants. Only strain 2009_45 was resistant to the third-generation cephalosporin cefotaxime, likely due to the presence of the *bla*_OXA-1_ gene. However, this resistance was not observed in strain 2009_38 which contained an almost identical antimicrobial resistance region, suggesting this gene is not expressed in this strain. Interestingly, both of these strains also showed phenotypic resistance to chloramphenicol despite only 2009_45 containing a complete copy of the *catA1* gene. The full repertoire of antibiotic resistance genes found in the 16 Sydney ST73 strains is presented in Figure 1.

**Table 1:**
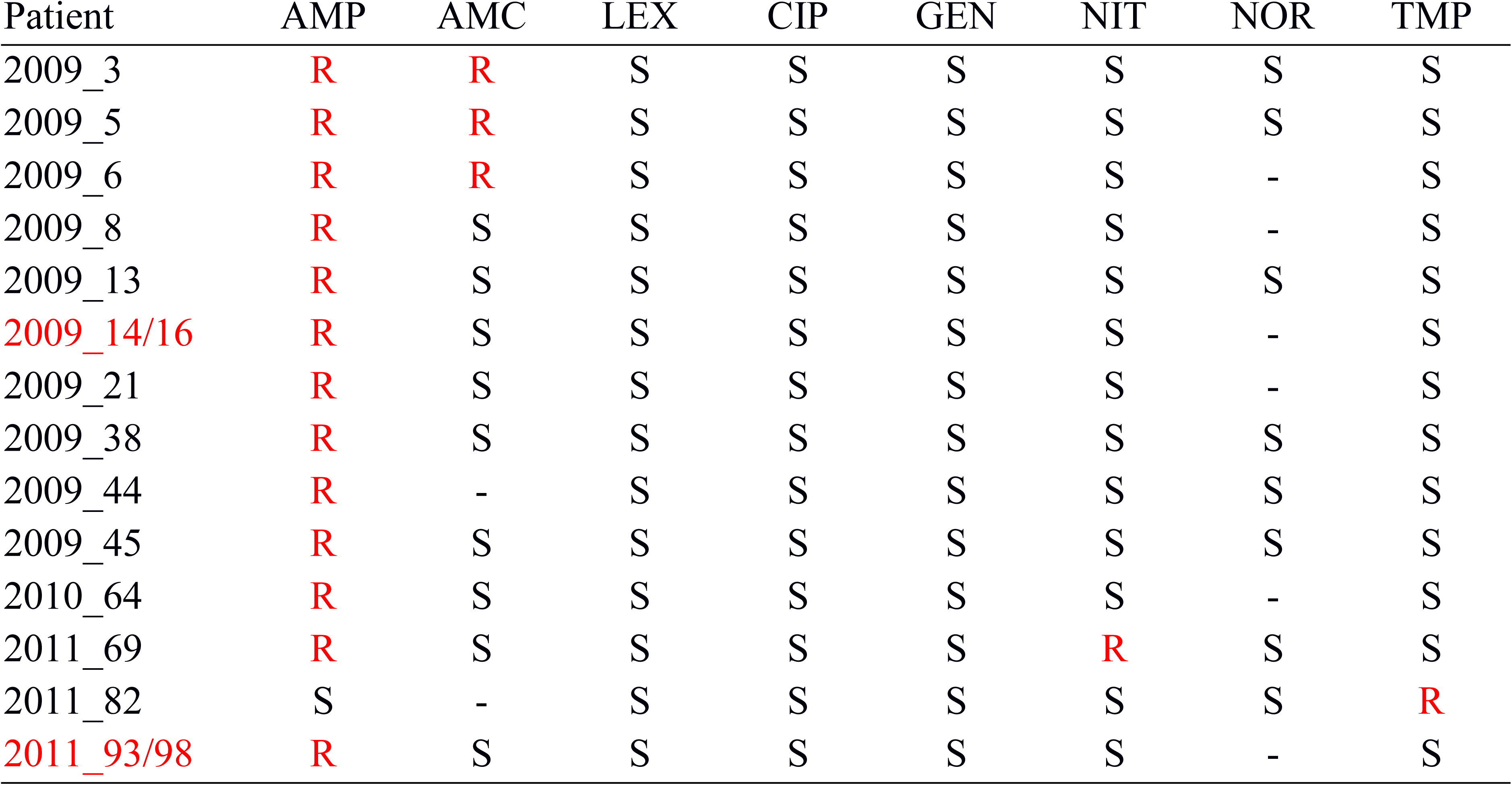
Phenotypic antimicrobial resistance

| Patient | AMP | AMC | LEX | CIP | GEN | NIT | NOR | TMP |
| --- | --- | --- | --- | --- | --- | --- | --- | --- |
| 2009_3 | R | R | S | S | S | S | S | S |
| 2009_5 | R | R | S | S | S | S | S | S |
| 2009_6 | R | R | S | S | S | S | - | S |
| 2009_8 | R | S | S | S | S | S | - | S |
| 2009_13 | R | S | S | S | S | S | S | S |
| 2009_14/16 | R | S | S | S | S | S | - | S |
| 2009_21 | R | S | S | S | S | S | - | S |
| 2009_38 | R | S | S | S | S | S | S | S |
| 2009_44 | R | - | S | S | S | S | S | S |
| 2009_45 | R | S | S | S | S | S | S | S |
| 2010_64 | R | S | S | S | S | S | - | S |
| 2011_69 | R | S | S | S | S | R | S | S |
| 2011_82 | S | - | S | S | S | S | S | R |
| 2011_93/98 | R | S | S | S | S | S | - | S |

### Structure of class 1 integrons in ST73 strains from Sydney

All locally sourced strains in this study, excepting 2011_82, were positive for a complete copy of the sulfonamide resistance gene *sul1*, a structural marker of the 3′-conserved segment (3′-CS) of class 1 integrons. Similarly, all strains contained the aminoglycoside resistance gene cassette *aadA1*. Strain 2011_82 was found to contain only a class 2 integron carrying the standard *dfrA1-sat2-aadA1* cassette array.

There were three class 1 integron-containing resistance regions represented within our collection (Fig. 3), all containing the same base structures with minor variations. The first structure was identified in 11 out of 15 class 1 integron-containing isolates (Fig. 3A). It consisted of an In-2 type class 1 integron with and *aadA1* gene cassette housed within an incomplete Tn*21* transposon, matching (99% sequence identity) to the sequence in the R100 plasmid identified in Japan in the 1950s from *Shigella flexneri* Accession NC_002134.1 [48]. However, our structure bears an IS*26*-mediated partial deletion of the Tn*21 tnpR* gene, which is a signature that has been reported previously twice within a uropathogenic *E. coli* strain from Australia, and in association with a different class 1 integron structure [49]. A Tn*3* transposon has inserted within the *mer* module of Tn*21* with partial deletion of *merA* and *merT* and complete deletion of *merC* and *merP*. The transposon is abutted downstream of *merR* by an inward facing IS*1* insertion element. One strain, 2009–64, housed this exact structure apart from the Tn*21 tnpM*, which appears to have been lost due to an IS*26*-mediated deletion event.

**Figure 3.**
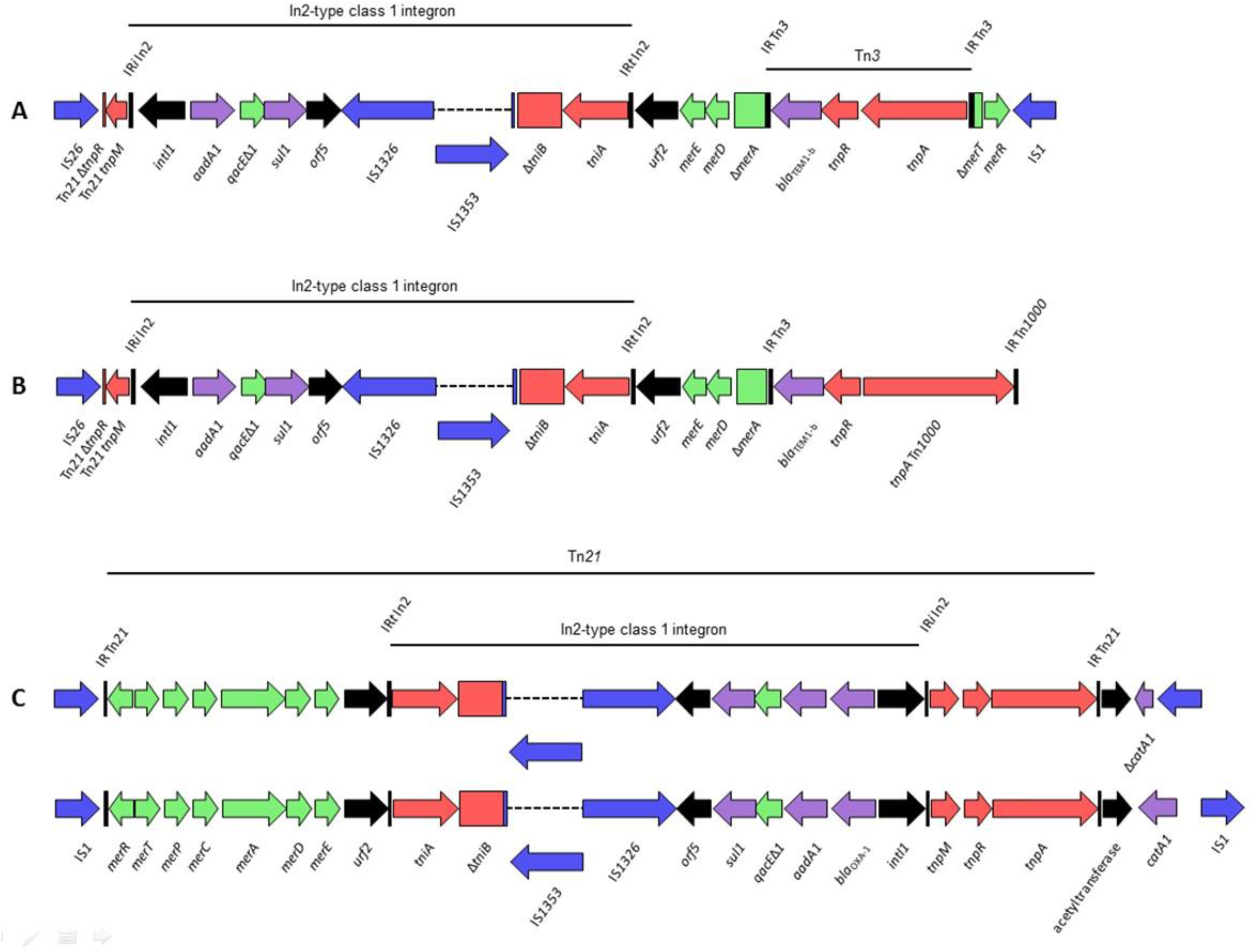
Schematic representations of integron structures found within this collection.

The complex resistance locus (CRL) shown in Figure 3B was identified in isolates 2009–6 and 2011–69, and shares homology with the structure in Figure 3A. It bore identical IS*26* and Tn*3* insertion points, with the only major difference being a crossover event where the standard Tn*3 tnpA* gene and terminal inverted repeat have been replaced by that of Tn*1000*, a transposon originally identified in a cosmid clone of a human DNA sequence in 1995 [50]. This signature was recently identified in the sequence of an unannotated plasmid of a *Salmonella enterica* serovar Typhi strain sequenced as part of a larger study of Typhi from typhoid-endemic regions of Asia and Africa (Accession LT904889.1). This is therefore the first report of this hybrid transposon and its presence in an *E. coli* isolated in Australia. Due to the nature of Illumina sequence technology we have no confirmed sequence information downstream of Tn*3*/Tn*1000*.

Structure 3 (Fig. 3C) shares homology to the previously discussed structures. However, this CRL, present in strains 2009–38 and 2009–45, has a *bla*OXA-1 gene cassette within the integron cassette array in addition to *aadA1*. Here, the Tn*21* transposon housing the class 1 integron is complete, with both the initial and terminal inverted repeats intact, and has an inward facing IS*1* flanking its *mer* end. There are two variants of this CRL in our collection, one of which contains a complete *catA1* gene downstream of Tn*21* followed by a second IS*1* element in the same orientation as the first, with the intergenic ORF identified as an acetyltransferase. This appears to be an established insertion event, with numerous reports in GenBank. In the second variant, the terminal IS*1* is inverted with consequent deletion of 476 bp of the *catA1* gene, forming a signature unique to isolates 2009-38 and 2009-45. This integron has been reported in its entirety in *Shigella dysenteriae* 1 plasmid p3099-85 (KT754164.1), *Salmonella enterica* serovar Typhimurium plasmid pUO-StVR2 (AM991977.1) and *Salmonella enterica* serovar Typhimurium strain T000240 (AP011957.1) Less than 10 SNPs were identified in comparative sequence alignments spanning the integron.

Sixty genomes from the SRA cohort returned adequate alignments to integron marker genes, with 25 of these appearing to have only the base class 1 integron with an *aadA1* cassette but no indication of a bordering Tn*21* transposon. Eight contained an *intI1* gene but no *aadA1*, suggesting the presence of a class 1 integron with a different cassette array. Sixteen contained *aadA1* but no class 1 integrase; this could indicate a deletion event or more likely the presence of *aadA1* in a class 2 integron context. Five genomes contained an unidentifiable integron structure, possibly variants of those described in the Sydney collection though it is impossible to say this definitively from read alignments against the abridged gene database used here.

Only three genomes; HVH_93_4–5851025, M0D1-EC6690 and MOD1-EC6783, contained all marker genes necessary to potentially contain integron C (Figure 3). However, within the SRA cohort, the presence of integrons A and B could not be confirmed.

All 16 strains of ST73 that were sourced from Sydney were shown to carry one or more plasmids (up to five) that ranged in size from 15 to 180 kb. Only one plasmid in each strain hybridized with the *intI1* probe (data not shown).

## Discussion

This study forms a part of wider global efforts to further understand the structure of disease-causing ST73 clones. Whole genome sequencing and maximum-likelihood phylogenetic analyses of these clones is providing important information on the community structure of ExPEC. Here we examined 16 ST73 isolates sourced from a single hospital and used sequence data sourced from the Sequence Read Archive to place these isolates into a broader global context and aid in identifying clonal lineages. Phylogenetic trees from this combined dataset, when overlaid with geographical and temporal data sourced from EnteroBase (data not shown), indicate that ST73 is globally disseminated in a manner similar to ST131 [51, 52] which is currently the most studied pandemic ExPEC lineage due to the frequency of CTX-M gene carriage. However, while ST131 tends to be relatively conserved in terms of core genome, ST73 appears more variable. Analysis of locally sequenced strains and comparison to globally-sourced reads from public databases can provide context which can allow the identification of outbreak clusters with more confidence than using total SNP counts alone and may help elucidate key outbreak groups and improve public health control of disease. This is valuable as the identification of clonal groups associated with outbreaks within larger bacterial populations remains a challenge.

Characterisation of molecular signatures can also assist in the identification of outbreaks as their transfer requires physical proximity of cells. CRL including integrons and transposons are common sites of genetic rearrangement and frequently carry unique molecular signatures due to insertion elements such as IS*26* (Dawes et al., 2010; Roy Chowdhury et al., 2015; McKinnon et al., 2018). While all class 1 integrons in the Sydney collection are not necessarily novel, there are IS-mediated signature deletions which do not appear to have been widely reported based on current literature. This suggests that these are local integron variants, an idea consistent with the lack of these structures in the global SRA cohort. The major representative class 1 integron described here has been reported in its entirety once within an Australian *E. coli* O2:K1:H7 ST95 strain isolated from a bloodstream infection in 2010 (unpublished data; GenBank accession CP021289.1). This integron also shares an IS*26*-mediated deletion of the Tn*21* resolvase gene *tnpR* with plasmid pUO-SeVR1 from a Spanish *Salmonella enterica* serovar Enteritidis strain sourced from a child with gastroenteritis [49, 53]. This is significant as this precise signature is likely the product of a single event. As such, a lateral transfer event is a likely explanation for the occurrence of this signature in disparate and geographically separate strains followed by changes in class 1 integron cassette content. It is likely that transfer of these integrons is being facilitated by IncF plasmids similar to pUO-SeVR1, as this is the major plasmid incompatibility type within our ST73 collection and our S1-PFGE data confirm that the class 1 integrons described here are plasmid-borne. Plasmids appear to increasingly play an important role in the mobilisation of drug resistance genes in ExPEC ST73, and their characterization relies heavily on the use of whole-genome sequencing (ideally long-read) and read-mapping technologies such as those described here.

Whole genome sequencing allows for the analysis of gene presence/absence in clinical isolates which will provide data on the importance of virulence genes in pathogenesis. The virulence profiles of strains sequenced in this study are consistent with other examinations of virulence in ST73 and in ExPEC more broadly. Genes encoding P fimbrial adhesins, the aerobactin siderophore *(iuc/iut*), and toxins hemolysin A and cytotoxic necrotizing factor 1 are not universally identified in worldwide ExPEC populations sourced from humans and animals [1, 54–56]. In previous work on ST73 isolates sourced from the UK, the prevalence of these genes/gene families was also non-universal; however, *hlyA* and *cnf1* showed substantially higher prevalence in ST73 compared with ExPEC-associated ST10, ST69 and ST95 [1]. In isolates sourced from Sydney, a relatively clear association could be identified between phylogenetic groups and virulence profiles. Further *in silico* categorization of virulence profiles using global ST73 reads would provide insight into virulence patterns/groups within ST73 and ExPEC, which could potentially lead to improved response, prevention and treatment of ExPEC linked-disease.

In endemic pathogens like *E. coli*, genetic comparisons of clonal group and mobile genetic element diversity can be difficult to perform with localized populations as high numbers of closely related isolates are required for robust SNP-phylogenetic analysis and this may require the long-term collection of bacterial isolates to isolate a sufficient number of representatives. Here we used Illumina sequencing combined with SNP-phylogenetic methods to identify at least eight distinct clonal lineages in a pilot sample of 16 ST73 isolates collected from a single hospital, indicating the wealth of diversity within ST73 population sourced from highly localised sampling over an extended period (Fig. 4). Contrastingly, the diversity of mobile elements within this cohort is much less profound. Only three resistance containing class 1 integron structures were identified, all were linked to plasmids, and all showed high structural similarity. Our study is an example of how genome sequencing can provide a depth of information not available with previous molecular epidemiology methodologies that are useful in the determination of outbreak groups among ST73 and the study of mobile genetic transfer in local populations.

**Figure 4.**
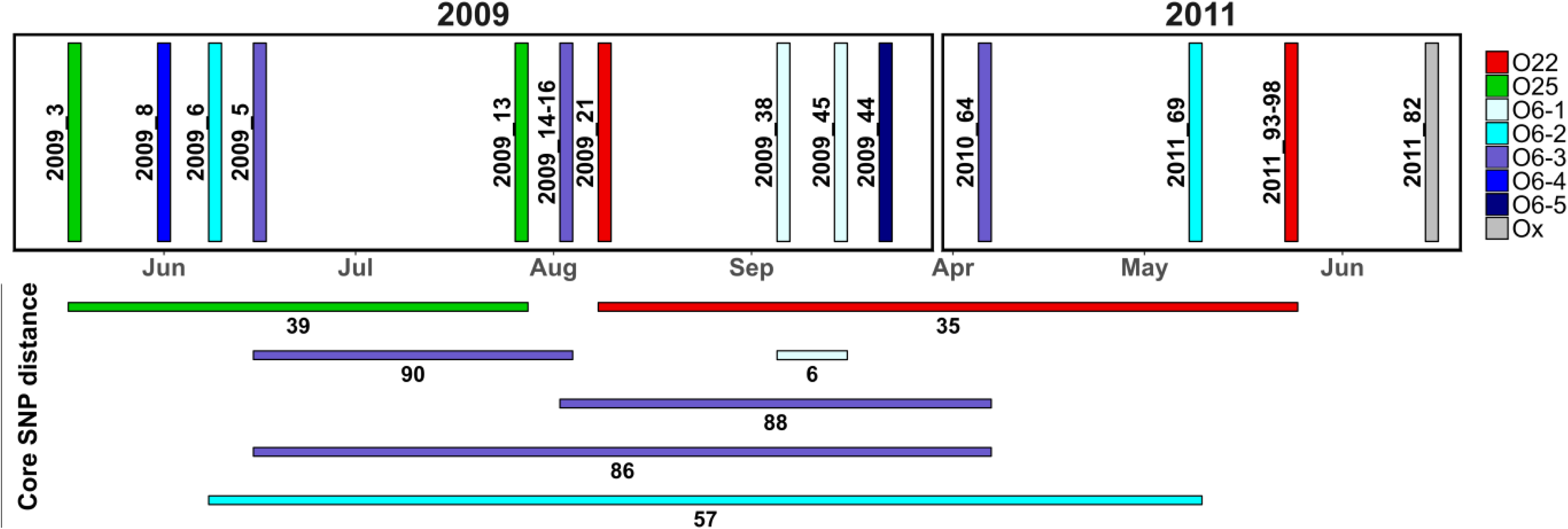
Epidemic curve for ST73 isolates sequenced in this study. Isolates are coloured to match groups identified by comparison to isolates from the Sequence Read Archive. Core SNP distances between samples of the same group are shown with coloured horizontal lines.

## Author Statements

DB performed the phylogenetic analysis, contributed to gene presence analyses, prepared most of the Figures, Tables and supplementary data and drafted multiple iterations of the manuscript. JM isolated genomic DNA, contributed to gene presence analyses, characterised structures of class 1 integrons of Sydney strains as well as generated the figure and contributed to writing the manuscript. ML constructed libraries and sequenced the Sydney ST73 strains. CV and JI performed the S1-PGFE and southern blot analyses and edited final drafts of the manuscript. AD assisted with data analysis. PRC initiated the collaboration with Sydney Adventist Hospital, screened isolates to create the strain collection included in this study and helped JM with analysis of the data presented. SD initiated and coordinated the project and drafted iterations of the final manuscript.

## Acknowledgements

We acknowledge the efforts of staff from the Sydney Adventist Hospital for providing the Sydney ST73 strains and associated metadata for this study.

## Ethical statement

Ethics clearance was not required.

## Conflicts of interest

There are no conflicts of interest to declare Abbreviations

**Abbreviations**
ST: Sequence type
ExPEC: Extraintestinal *Escherichia coli*
SNP: Single nucleotide polymorphism
S1-PFGE: S1 Nuclease - Pulsed Field Gel Electrophoresis
CS: Conserved segments
UPEC: Uropathogenic *E. coli*
NMEC: Neonatal meningitis-causing *E. coli*
APEC: Avian pathogenic *E. coli*
UTI: Urinary tract infection
VAG: Virulence-associated gene
MDR: Multiple drug resistant
SAN: Sydney Adventist Hospital
LB: Luria-Bertani
AMP: Ampicillin
AMC: Amoxicillin-clavulanic acid
LEX: Cephalexin
CIP: Ciprofloxacin
GEN: Gentamicin
NIT: Nitrofurantoin
NOR: Norfloxacin
TMP: Trimethoprim

## Supplemental material

1. All sequencing reads and assemblies for isolates sequenced in this study have been submitted to the ENA Sequence Read Archive and GenBank, respectively. GenBank, SRA accession numbers and URLs are included in Table S1.
2. Scripts used for the analysis of SNP phylogeny have been deposited in Github; (URL - https://github.com/bogemad/snp_phylogeny)
3. Figure S1 has been deposited in Figshare; DOI: 10.6084/m9.figshare.5477449 (URL – https://figshare.com/s/b720dcd41ca1c11fdae2)
4. Figure S2 has been deposited in Figshare; DOI: 10.6084/m9.figshare.5477485 (URL – https://figshare.com/s/0835e6174bf9f026ff61)
5. Table S1 has been deposited in Figshare; DOI: 10.6084/m9.figshare.5477461 (URL – https://figshare.com/s/e6ea61f9dde79e9b18dd)
6. Table S2 has been deposited in Figshare; DOI: 10.6084/m9.figshare.5477464 (URL – https://figshare.com/s/70de292aa98e806dd05f)
7. Table S3 has been deposited in Figshare; DOI: 10.6084/m9.figshare.5477467 (URL – https://figshare.com/s/ddf02248b7f2d6f6ebe6)
8. Table S4 has been deposited in Figshare; DOI: 10.6084/m9.figshare.5477473 (URL – https://figshare.com/s/b91bbc880eb31e5f90fc)
9. Table S5 has been deposited in Figshare; DOI: 10.6084/m9.figshare.5477476 (URL – https://figshare.com/s/3b8be84c33225bb32d78)
10. Table S6 has been deposited in Figshare; DOI: 10.6084/m9.figshare.5477479 (URL – https://figshare.com/s/7ba306327343fdeca167)
11. Additional supplemental vcf files of total and core SNPs identified from the whole genome phylogeny of 226 isolates have been deposited in Figshare; Core SNP - DOI: 10.6084/m9.figshare.5477458 (URL – https://figshare.com/s/e033c384eca1eab880e7); Total SNP - DOI: 10.6084/m9.figshare.5477455 (URL – https://figshare.com/s/fd0233d6cca641d44bee)

